# Neonatal Mouse Ovary Culture: An In Vitro Model for Studying Primordial Follicle Regulation

**DOI:** 10.1101/2025.08.19.671127

**Authors:** Edgar A Diaz Miranda, Grace Anne Dyer, Faith Wilson, Mariel English, Taylor VanDeVoorde, Lei Lei

## Abstract

In mammalian females, primordial follicles form during fetal ovarian development and serve as the only source to sustain adult ovarian function. Mechanisms underlying how primordial follicles assemble, maintain dormancy, activate for follicular development, and undergo cell death are important for understanding ovarian physiology and pathological conditions. Here, we demonstrate a protocol of culturing postnatal mouse ovaries on membrane inserts - an approach allowing culture, pharmaceutical treatment, and live-imaging of intact ovaries for up to 10 days depending on the developmental stage of the ovary. The change of culture conditions can be done by transferring inserts containing cultured ovaries between wells on a plate, avoiding physical interference with tissues during culture. In this experiment, we use postnatal day 5 (P5) CD1 mouse ovary culture as an example. P5 ovaries were isolated and placed on a 12 mm insert in a 24-well plate. Each ovary was separated within a droplet of DMEM/F12 medium supplemented with 10% FBS, 3 mg/ml BSA, 10 mIU/ml FSH, and Gibco™ Antibiotic-Antimycotic, and gently stabilized to the membrane insert. The medium was changed every two days, and the culture was maintained for five days. Following the culture, ovaries were fixed in 4% paraformaldehyde for two hours and processed for whole-mount antibody staining. Primordial follicles were visualized using confocal microscopy with anti-DDX4 antibody, allowing for the analysis of oocyte number and morphology. We showed that the number of primordial follicles in each ovary was significantly affected by whether tissues were properly placed on the membrane insert. Difference in the number of ovaries on each insert may contribute to non-biological variations and should be avoided.

## Introduction

Oogenesis is a highly regulated and complex process that ends in the production of mature oocytes. In mammals, oogenesis initiates in fetal ovaries, where primordial germ cells (PGCs) differentiate into primary oocytes. Each primary oocyte is further enclosed by a layer of squamous pregranulosa cells, forming a primordial follicle. Most of the primordial follicles become dormant after follicle formation and serve as the ovarian reserve^1-3^. In the adult ovary, periodical activation of primordial follicles to undergo follicular development (i.e. folliculogenesis) is an essential process for sustaining mature oocyte and ovarian steroid hormone production^4,5^.

The process of oogenesis is highly conserved in mammals, making mice an ideal model for studying mammalian oogenesis. In mice, primordial follicle formation completes around postnatal day 4 (P4). A small proportion of primordial follicles undergo follicular development immediately after follicle formation. In the developing follicle, the oocyte grows in size due to increased organogenesis and mRNA and protein synthesis^4^; squamous pregranulosa cells transition to cuboidal granulosa cells and become proliferative^6^. These developing follicles grow to ovulatory stage around day 21, when the female mice reach puberty. The follicular development in postnatal ovaries is often referred to as the ‘first-wave folliculogenesis’. Because first-wave folliculogenesis well recapitulates the process of follicle development, postnatal mouse ovaries serve as an ideal model for studying primordial follicle regulation and ovarian folliculogenesis^7^.

The relatively small tissue size of postnatal mouse ovaries makes whole ovary culture feasible. It provides a highly effective tool for studying ovary and follicle development within an intact ovary, since the culture maintains of ovarian physical and physiological microenvironments while preventing any interference from other tissues. This approach allows us to conduct experiments that are often infeasible in in vivo models. Examples include live imaging of ovary development, time-controlled multi-drug treatments, and analyzing ovarian secretory activity through protein profiling of the culture media. This approach can be utilized to study the effect of secretory factors without direct cell-cell interaction through co-culturing experiments, in which different types of tissues can be placed on separate membrane inserts in the shared media.

In this article, we introduce an approach of postnatal mouse ovary culture on the membrane insert, demonstrating techniques of 1) neonatal mouse ovary dissection, 2) neonatal mouse ovary culture, 3) media change during culture, 4) tissue fixation and whole mount antibody staining, and 5) tissue imaging and follicle quantification. Our results showed that ovary number per insert and ovary location in the insert during culture caused difference in primordial follicle numbers, emphasizing the importance of keeping the culture setup consistent to avoid non-biological experimental variations.

### Protocol

1. ***Ethics and mice*** CD-1 mice were purchased from Charles River Laboratories. Male and female mice were housed at 1:1 ratio as breeding pairs. The date of birth of new pups was considered postnatal day 0. All animal experiments were approved by the Institutional Animal Care & Use Committee (IACUC) at the University of Missouri (protocol number: 36647).
2. ***Reagents and Culture Media***
  1. All reagents used in this study are listed in **Supplementary Table 1**. Ensure to prepare all materials and media before ovary culture.
  2. Culture media: DMEM/F-12 media supplemented with 3 mg/ml bovine serum albumin (BSA), 10% fetal bovine serum (FBS) and 10 mIU/ml follicular stimulating hormone (FSH). Note: The media should be prepared freshly by adding the BSA (30 mg/ml), FBS, and FSH stock (10 IU/ml) to DMEM/F-12 media and equilibrate for about 30 minutes in a 5% CO2 incubator.
3. ***Ovary culture***
  1. Euthanize P5 female mice by decapitation. Remove the ovaries and the attached bursa tissue using surgical scissors and forceps. Place the ovaries in a 60 mm petri dish with pre-chilled Dulbecco’s phosphate buffered saline (DPBS) containing Gibco™ Antibiotic-Antimycotic.
  2. Using a pair of insulin needles, dissect and remove bursa tissue from the ovary under a stereoscope. Transfer the ovaries to a 30 mm petri dish with the culture media and place the dish in a CO_2_ incubator.
  3. Under the laminar flow hood, prepare the plate by placing 400 μl of media and an insert in each well (**Figure 1A, A’**). Pre-soak the insert by transferring ~200 μl of media from the well to the insert and then transferring the media back to the well. For the culture with 24 hr Doxorubicin (Dox) treatment, prepare media containing Dox at 0.1 μg/ml concentration freshly by diluting Dox stock (10 mg/ml) with the media.
  4. Under a stereoscope, transfer up to three ovaries to an insert using a 1 ml pipette. Try to minimize the amount of media going to the insert during transfer and immediately remove any excess media in the insert. Use a 200 μl pipette with media from the well to place the ovaries properly. Make sure that proper space is left between the ovaries and that they are well placed/stabilized on the membrane insert (**Figure 1B**), rather than remaining suspended in the medium (**Figure 1C**).
  5. In a 5-day culture, change media every two days by removing 200 μl media (half of the total amount) from the well and adding 250 μl fresh culture media back to the well. We add a little more fresh media because there is usually a slight loss of media during culture due to evaporation.
  6. For the culture with 24 hr drug treatment, the insert with ovaries are transferred to a new well with fresh culture media after the initial 24 hrs culture using the following steps: take the insert with ovaries using a pair of forceps, rinse the bottom of the insert briefly in a 30 mm dish with culture media, gently tap the bottom of the insert on an empty 30 mm dish to remove the excessive media, and place the insert into the well with fresh culture media.
4. ***Fixation and whole mount immunostaining***
  1. For tissue fixation, take the insert out of the plate using a pair of forceps and place the insert in a petri dish with DPBS. Add ~500 μl DPBS to the insert to resuspend the cultured ovaries using a 1ml pipette. **Note**: The end of the pipette tip may need to be trimmed wider if the ovaries are too large to go through the it.
  2. Using a 1 ml pipette, transfer the ovaries from the insert to the dish with DPBS for a quick rinse before transferring the ovaries to a 1.5 ml Eppendorf tube. Remove DPBS from the tube and add 500 μl 4% paraformaldehyde. Fix the ovaries at 4 °C for 2 hrs.
  3. Wash the fixed ovaries in PBST_2_ (PBS, 0.1%Tween, 0.5% Triton) at least three times, 30 mins each on a rotating shaker. Note: Longer washes improve antibody penetration.
  4. Incubate the ovaries with anti-DDX4 (Cell signaling-Cat#8761) in 100 μl antibody dilution buffer (1% BSA, 10% Donkey serum, 0.1 M glycine, 0.1% Tween and 0.5% Triton in PBS) at a dilution of 1/400 at 4 °C overnight.
  5. On the next day, remove the primary antibody and add 1ml PBST2 to the tube. Wash three times (at least 30 min each) in PBST_2_. **Note**: This can be a stopping point, the tissues can remain in PBST_2_ up to three days after primary antibody incubation.
  6. Dilute the secondary antibody (Donkey anti-rabbit AF488, cat# 711-546-152) at 1/500 dilution in PBST_2_. Incubate the tissue with at least 100 μl of the diluted secondary antibody at 4 °C overnight.
  7. On the next day, wash the tissues extensively three times (at least 30 min each) and incubate in DAPI for 30 min at room temperature at a dilution of 1/1000 in PBS.
  8. Wash ovaries in PBST_2_ once and mount them on a microscopic slide with a spacer (Grace Bio-Labs SecureSeal^TM^ imaging spacer). Transfer the ovaries to the center of the spacer and remove excess PBST_2_. Add about 10 μl of the antifade mounting medium (VectaShield®), and place a cover slip on the top of the spacer. Seal the edges of the cover slip with clear nail polish for long term storage. **Note**: Before imaging the slides, ensure nail polish is completely dry and there is no mounting medium on the outside of the cover slip. These reagents may cause damage to the objectives of the microscope.
  9. Image the ovaries with a confocal microscope (Leica TCS SP8 system) at 10x magnification, with a Z-step size of 10 μm. Adjust settings and use the Z-compensation (excitation gain) to image the entire ovary.
5. ***Follicle quantification***
  1. The images acquired by confocal can be viewed and analyzed in 3D using Imaris software (https://imaris.oxinst.com/) (**Figure 2 A and B**) or Image J software (https://imagej.net/ij/)^8^.
  2. To use Image J for follicle quantification, open the images and use ‘image/transform/rotate’ tool to place the ovary symmetrically in the image.
  3. Use the ‘analyze/tools/grid’ tool to divide the ovary into grids (**Figure 2C**).
  4. Use the ‘plugins/analyze/cell counter’ tool to mark all the grids covering the ovary (**Figure 2C’**). Of these grids, mark every other grid in each row for follicle quantification (**Figure 2C’’**, grids marked by 8). Grids on the edge of the ovary that have less than 10 oocytes are not included for follicle quantification.
  5. Primordial follicles are recognized by DDX4-positive oocytes at the size of about 20 μm in diameter surrounded by squamous somatic cells. Developing follicles are recognized by DDX4-positive oocytes that are greater than 25 μm in diameter and surrounded by cuboidal granulosa cells.
  6. To count the follicles in each grid marked by 8 in Figure 2C’’, go over optical sections/stacks in the grid and mark primordial and developing follicles using cell counter tool, respectively. Ensure not to double count primordial follicles on two consecutive optical sections. Since half of the ovary was counted, the number of follicles in the entire ovary (nf) equals the average number of follicles per grid (nfgrid) multiplied by the number of grids of the ovary (ngrids): *nf* = *nfgrid* ×*ngrids*.
6. ***Statistical analysis*** Statistical analyses were performed and graphs were generated using Prism 10.2.0 software (GraphPad Software, Version 10, Inc., CA). Differences among groups were evaluated using one-way analysis of variance (ANOVA) followed by Tukey’s multiple comparison test. Statistical significance was considered at p < 0.05. Data is presented as the mean ± standard error (SE).

**Figure 1.**
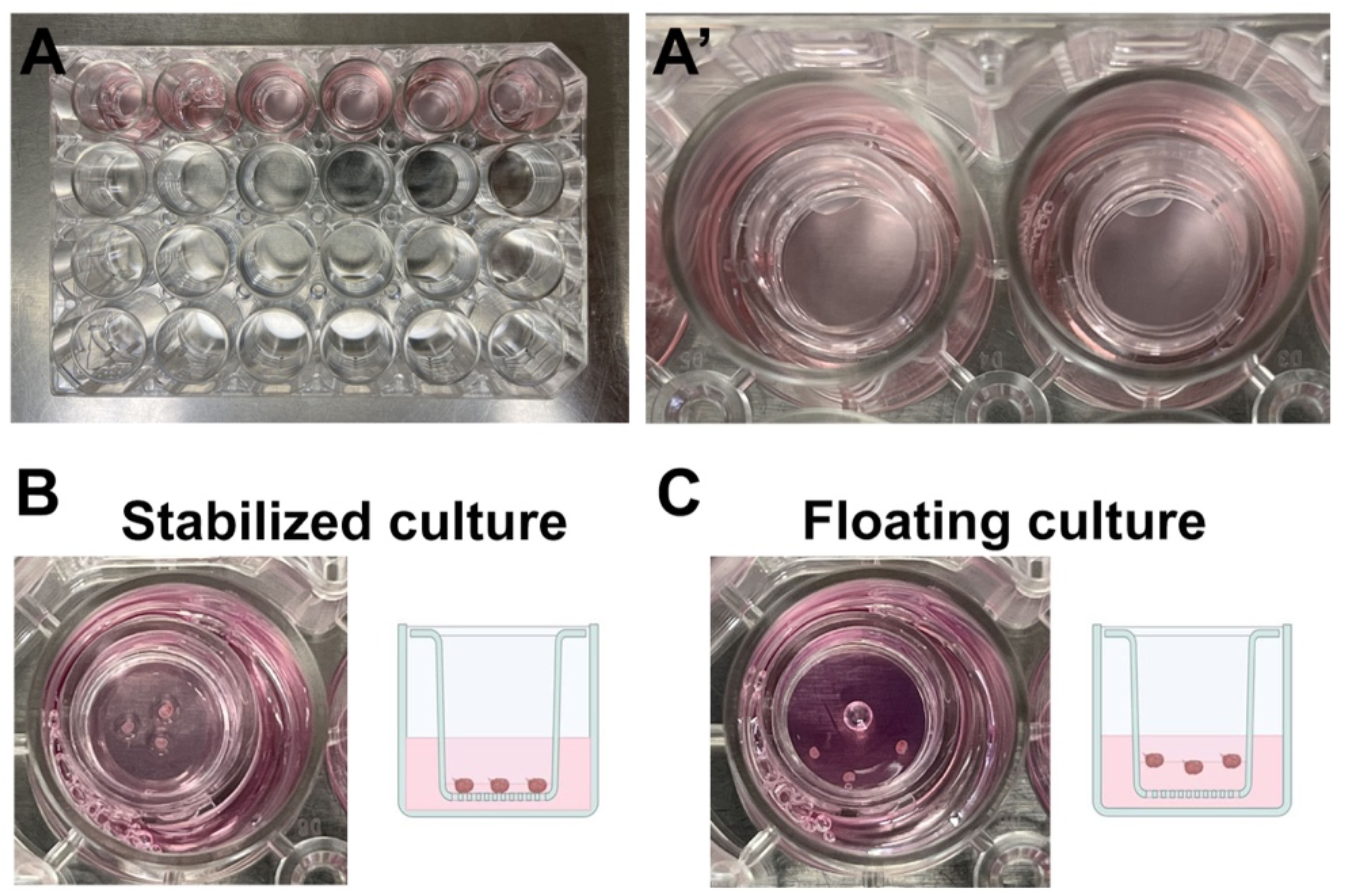
Representative images of mouse ovarian culture setup. (A-A′) Inserts placed into wells containing 400 µL of medium. (B-C) Wells showing two experimental groups: three ovaries stabilized and three ovaries floating. Images (B-C) were created with BioRender.com.

**Figure 2.**
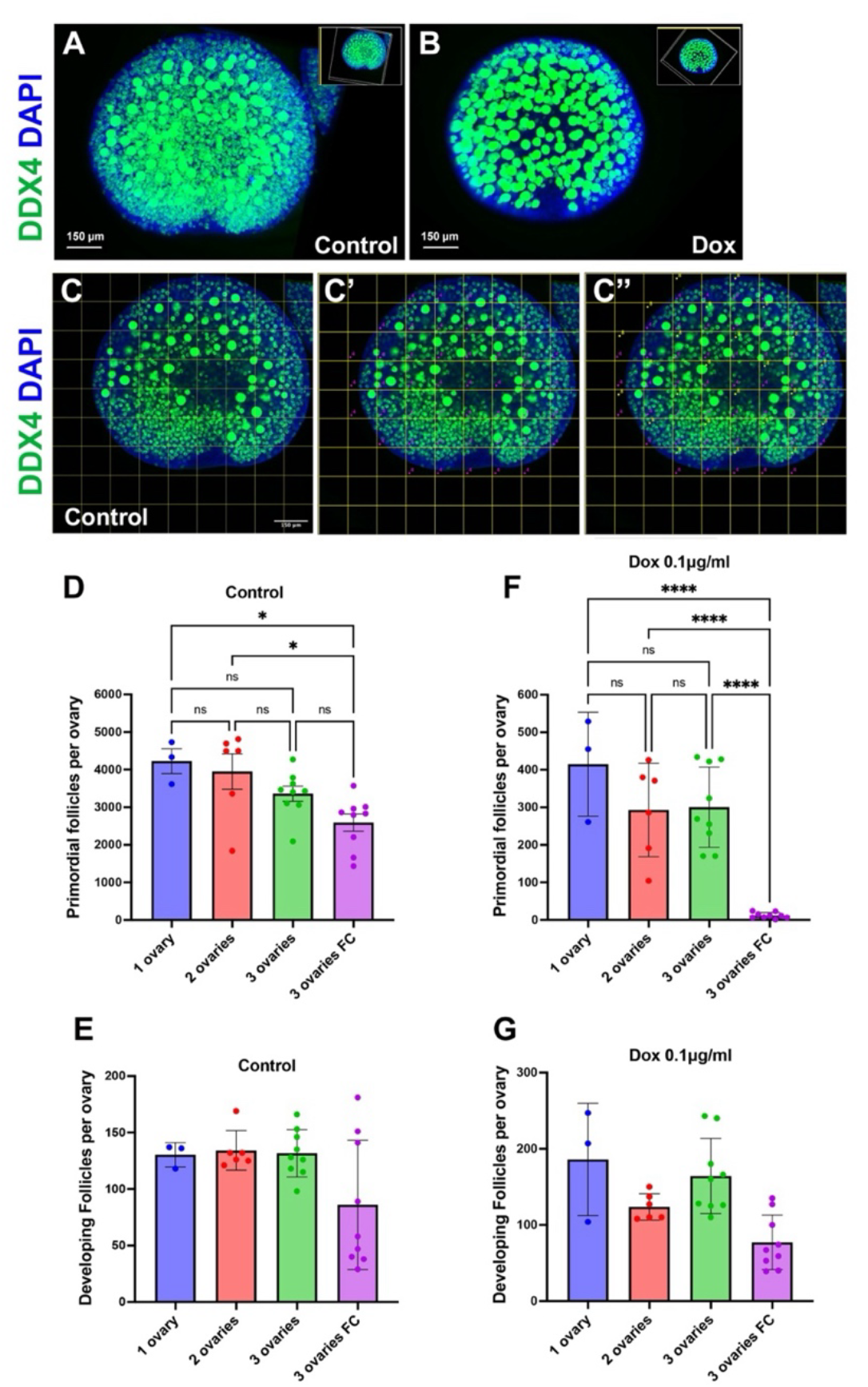
Follicle quantification and representative results. (A-B) 3D images from Imaris showing control and Dox-treated ovaries. (C-C′-C′′) Ovarian quantification using the Grids tool in ImageJ, indicating the total number of grids selected in the ovary (marked with the number 4) and the number of grids counted (marked with the number 8). (D-E) Numbers of primordial and developing follicles per ovary after 5 days of culture under different conditions (1 ovary, 2 ovaries, or 3 ovaries in the stabilized condition, and 3 ovaries in the floating condition). (F-G) Numbers of primordial and developing follicles per ovary after culture with Dox under different culture conditions. Data are presented as mean ± standard error (S.E.). *p < 0.05; ****p < 0.0001.

### Representative results

In this article, through demonstrating the approaches of neonatal ovary culture, whole mount antibody staining, confocal imaging and follicle quantification, we assessed whether the number and the position of the ovaries in the insert affect culture outcome, in particular, follicle numbers in the ovary. We cultured ovaries with the following set up: one, two, or three ovaries per insert for stabilized culture, and three ovaries per insert for floating culture. The ovaries were cultured under control condition or drug-treated condition (24 hr treatment with Dox at 0.1 μg/ml). The ovaries of both control and Dox-treated groups were switched to fresh culture media after 24 hr culture.

We found that for the ovaries cultured in control media with the stabilized culture approach, although on average there were more primordial follicles in the one-ovary culture group, and less primordial follicles in the three-ovary group, there was no significant difference between the three groups. The three-ovary group cultured using floating culture approach had the least number of primordial follicles among the four culture conditions. However, no significant difference was observed between three-ovary cultured in stabilized and floating approaches (**Figure 2D**). Ovaries cultured in the floating approach also had the least number of developing follicles. Ovaries cultured using the stabilized approach had similar number of developing follicles regardless the number of ovaries cultured in each insert. No significant difference in the numbers of developing follicles was observed among the four groups (**Figure 2E**).

For the ovaries cultured with 24 hr Dox treatment, among the three groups with stabilized culture, the one-ovary group had the most primordial follicles, although no significant difference was observed among the three groups. Strikingly, three ovaries cultured in the floating approach had significantly less primordial follicles compared with ovaries cultured in the stabilized approach regardless of the number of ovaries in the insert (**Figure 2F**). There was no significant difference when comparing developing follicles among the four culture groups. On average, three ovaries cultured in the floating approach had the least number of developing follicles (**Figure 2G**).

## Discussion

In this study, we demonstrated a neonatal mouse ovary culture system, which can sustain primordial and early developing follicles. Our results indicate that it is important to keep the ovaries cultured in the stabilized approach, which provides consistent physical support to the ovary. Ovaries cultured in the floating approach had an increased primordial follicle loss that may be primarily due to cell death in these follicles, as we did not observe an increased number of developing follicles in the ovaries cultured with the floating approach. The difference in primordial follicle numbers between the stabilized culture approach and the floating culture approach may cause non-biological variations in experiment results. Keeping ovaries stabilized during the culture is particularly important when conducting drug treatment, since significantly less primordial follicles were found in the Dox-treated ovaries cultured in the floating approach. We observed that when ovaries were cultured in the stabilized approach, the number of ovaries cultured in each insert caused a slight difference in average primordial follicle numbers. Thus, it is important to keep ovary number per well consistent throughout the study to avoid non-biological variations. In summary, neonatal mouse ovary culture is an effective approach for studying primordial follicles, and it is essential to keep the culture conditions as consistent as possible.

## References

1. Pepling ME. From primordial germ cell to primordial follicle: mammalian female germ cell development. Genesis. Dec 2006;44(12):622–32. doi:10.1002/dvg.20258

2. Ikami K, Nuzhat N, Lei L. Organelle transport during mouse oocyte differentiation in germline cysts. Curr Opin Cell Biol. Feb 2017;44:14–19. doi:10.1016/j.ceb.2016.12.002

3. Pepling ME. Follicular assembly: mechanisms of action. Reproduction. Feb 2012;143(2):139–49. doi:10.1530/REP-11-0299

4. Lintern-Moore S, Moore GP. The initiation of follicle and oocyte growth in the mouse ovary. Biol Reprod. May 1979;20(4):773–8. doi:10.1095/biolreprod20.4.773

5. Picton HM. Activation of follicle development: the primordial follicle. Theriogenology. Apr 1 2001;55(6):1193–210. doi:10.1016/s0093-691x(01)00478-2

6. Binelli M, Murphy BD. Coordinated regulation of follicle development by germ and somatic cells. Reprod Fertil Dev. 2010;22(1):1–12. doi:10.1071/rd09218

7. Xu M, Kreeger PK, Shea LD, Woodruff TK. Tissue-engineered follicles produce live, fertile offspring. Tissue Eng. Oct 2006;12(10):2739–46. doi:10.1089/ten.2006.12.2739

8. Schneider CA, Rasband WS, Eliceiri KW. NIH Image to ImageJ: 25 years of image analysis. Nat Methods. Jul 2012;9(7):671–5. doi:10.1038/nmeth.2089

